# Vaccine blunts fentanyl potency in male rhesus monkeys

**DOI:** 10.1101/580100

**Authors:** Rebekah D. Tenney, Steven Blake, Paul T. Bremer, Bin Zhou, Candy S. Hwang, Justin L. Poklis, Kim D. Janda, Matthew L. Banks

## Abstract

One proposed factor contributing to the increased opioid overdose deaths is the increased frequency of synthetic opioids, including fentanyl and fentanyl analogs. A treatment strategy currently under development to address the ongoing opioid crisis is immunopharmacotherapies or opioid-targeted vaccines. The present study determined the effectiveness and selectivity of a fentanyl-tetanus toxoid conjugate vaccine to alter the behavioral effects of fentanyl and a structurally dissimilar mu-opioid agonist oxycodone in male rhesus monkeys (n=3-4). Fentanyl and oxycodone produced dose-dependent suppression of behavior in an assay of schedule-controlled responding and antinociception in an assay of thermal nociception (50°C). Acute naltrexone (0.032 mg/kg) produced an approximate 10-fold potency shift for fentanyl to decrease operant responding. The fentanyl vaccine was administered at weeks 0, 2, 4, 9, 19, and 44 and fentanyl or oxycodone potencies in both behavioral assays were redetermined over the course of 49 weeks. The vaccine significantly and selectively shifted fentanyl potency at least 10-fold in both assays at several time points over the entire experimental period. Mid-point titer levels were significantly correlated with fentanyl antinociceptive potency shifts. Antibody affinity for fentanyl as measured by a competitive binding assay increased over time to around 3-4 nM. The fentanyl vaccine also significantly increased fentanyl plasma levels approximately 6-fold consistent with the hypothesis that the vaccine sequesters fentanyl in the blood. Overall, these results support the continued development and evaluation of this fentanyl vaccine to address the ongoing opioid crisis.

**Highlights:**

- Vaccine blunted fentanyl rate-suppression potency ∼ 10-fold
- Vaccine blunted fentanyl antinociceptive potency ∼25-fold
- Fentanyl vaccine was as effective as acute 0.032 mg/kg naltrexone
- Vaccine was selective for fentanyl and not oxycodone
- Antibody immune response ∼ 3 nM affinity for fentanyl

## 1.0 INTRODUCTION

The rates of fatal and non-fatal overdoses attributed to mu-opioid receptor (MOR) agonists have significantly increased every year since 2006 (O’Donnell et al., 2017). A recent Center for Disease Control report found that the synthetic MOR agonist fentanyl was detected in 56% of all reported overdose deaths (O’Donnell et al., 2017). The source of fentanyl driving the current opioid crisis is not originally from diverted prescription products, but primarily manufactured in Asian laboratories and trafficked into the United States where it is mixed with heroin or other MOR agonists (Ciccarone, 2017). Currently, the only Food and Drug Administration (FDA)-approved treatment for opioid overdose is the opioid antagonist naloxone. Although naloxone is effective, it has several limitations. First, the half-life of naloxone is approximately 30 min and may be shorter than most illicit and prescription opioids (for review, see (Kim and Nelson, 2015; Ryan and Dunne, 2018)). Second, naloxone will precipitate somatic withdrawal signs in opioid-dependent individuals (van Dorp et al., 2007). In summary, despite clinically available treatment options for opioid overdose, overdose deaths continue to occur at an increasing rate.

Recently, the National Institutes of Health have outlined several scientific areas of interest to strategically focus research efforts for developing novel treatment strategies to address the ongoing opioid crisis. One proposed strategic area is the development of opioid-targeted conjugate vaccines or immunopharmacotherapies (Baehr and Pravetoni, 2019; Bremer and Janda, 2017; Volkow and Collins, 2017). A conjugate vaccine consists of three components: a hapten (i.e., opioid analog), an immunogenic carrier protein to stimulate an immune response (e.g., tetanus toxoid, TT), and adjuvant(s) to boost the immune system (Bremer and Janda, 2017). These vaccines stimulate the immune system to produce high-affinity antibodies specifically against the targeted opioid. Upon exposure to the targeted opioid, antibodies in the blood bind the opioid, and the resulting antibody-opioid complex is too large to cross the blood brain barrier and cannot activate central opioid receptors that mediate either abuse-related effects or respiratory function. Potential advantages of conjugate vaccines include high selectivity towards the targeted opioid and a longer duration of action mitigating the need for repeated dosing (Baehr and Pravetoni, 2019; Banks et al., 2018).

Preclinical research is a critical component in the development of novel therapeutics to address the opioid crisis. Fentanyl vaccines have been developed and evaluated in both mice and rats (Bremer et al., 2016; Raleigh et al., 2019; Torten et al., 1975). For example, the fentanyl-tetanus toxoid (TT) conjugate vaccine used in the present study produced a 33-fold antinociceptive potency shift for fentanyl and a 9-fold shift antinociceptive potency for α-methylfentanyl in mice (Bremer et al., 2016). In addition, a fentanyl-keyhole limpet hemocyanin (KLH) conjugate vaccine produced a 5.4-fold antinociceptive potency shift for fentanyl in rats (Raleigh et al., 2019). Overall, these results in rodents support the continued evaluation of fentanyl vaccines in higher order species.

The aim of the present study was to examine the effectiveness and selectivity of a fentanyl-TT conjugate vaccine in rhesus monkeys. Vaccine effectiveness was evaluated on two behavioral endpoints and vaccine selectivity was compared to the structurally dissimilar and clinically prescribed MOR agonist oxycodone. Warm water tail-withdrawal was utilized to allow comparisons to previous rodent studies and schedule-controlled responding was utilized to assess MOR agonist potency shifts (Bremer et al., 2017; Butelman et al., 1996; Negus et al., 2003; Negus et al., 1993). Fentanyl vaccine effectiveness was determined in nonhuman primates for two mains reasons. First, immunological factors related to both total B-cell and T-cell counts are more similar between rhesus monkeys and humans than rodents and humans. (Caldwell et al., 2016; Vaccari and Franchini, 2010). Second, pharmacodynamic considerations related to MOR density and receptor distribution and pharmacokinetic considerations in opioid metabolism provide further support for the evaluation of candidate therapeutics in nonhuman primates as part of the drug development process (Weerts et al., 2007). Fentanyl vaccine effectiveness was compared to acute naltrexone treatment in the assay of schedule-controlled responding. We have previously reported that 0.032 mg/kg intramuscular (IM) naltrexone produced an approximate 9-fold shift in fentanyl antinociceptive potency (Cornelissen et al., 2018). Furthermore, human laboratory studies and clinical trials have established the minimally effective clinical naltrexone dose results in an 8-fold opioid potency shift (Bigelow et al., 2012; Comer et al., 2006; Sullivan et al., 2006).

## 2.0 METHODS

### 2.1 Subjects

Four adult male rhesus monkeys (*Macaca mulatta*) weighing between 10-14 kg served as subjects. All subjects had extensive drug and experimental histories, including opioids. Subjects were individually housed in stainless steel chambers that also served as the experimental chambers. Water was available ad lib. The primary food diet (Teklad Global Diet, 2050, 20% Protein Primate Diet) was given daily after the procedure and supplemented daily with fruits and vegetables. In addition, subjects could earn food pellets (TestDiet, Grain-Based Non-Human Primate Tablet) during the behavioral session. Housing rooms were maintained a 12-hour light cycle (6:00 AM to 6:00 PM). Animal maintenance and research were conducted in accordance with the 2011 guidelines promulgated by the National Institutes of Health Committee on Laboratory Animal Resources. The facility was licensed by the United States Department of Agriculture and accredited by AAALAC International. Both research and enrichment protocols were approved by the Virginia Commonwealth University Animal Care and Use Committee. Monkeys had visual, auditory, and olfactory contact with other monkeys throughout the study. Monkeys also had access to mirrors, television, puzzle feeders, chew toys, coconuts, and birch sticks as additional environmental enrichment.

### 2.2 Schedule-controlled responding procedure

Experiments were conducted in the housing chamber which also served as the experimental chamber as previously described (Banks et al., 2010; Bremer et al., 2017). Briefly, operant response panels were mounted daily on the front of each chamber. Each panel consisted of three-square translucent keys and a pellet dispenser (ENV-203-1000, Med Associates, St. Albans, VA, USA) that delivered 1-g banana-flavored pellets into a food receptacle beneath the operant panel. Operant panels and experimental parameters were controlled by a MED-PC interface and an IBM compatible computer. Custom programming in MEDSTATE Notation (MED Associates) was utilized.

The schedule-controlled responding procedure lasted 75 minutes in duration and consisted of five 15-minute cycle. Each cycle contained two components: a 10-min timeout period followed by a 5-min response period. During the response period, the right key was illuminated red. Subjects could respond on the key under a fixed-ratio 30 (FR30) schedule of reinforcement and receive a maximum of 10 pellets per cycle. If the maximum number of pellets was earned before the 5-min response period elapsed, the light turned off. Responding in the absence of an illuminated key had no programmed consequences. Experimental sessions were conducted 5 days per week. Training sessions occurred on Mondays, Wednesdays and Thursdays and test sessions occurred on Tuesdays and Fridays. Training sessions included either no injection or a saline injection at the beginning of each 10-min timeout period. All monkeys were trained until rates of responding were consistently ≥ 1.0 response/s before transitioning to test sessions.

Initially, dose-effect functions were determined for fentanyl (0.001-0.032 mg/kg) and oxycodone (0.0032-1.0 mg/kg), and each drug was tested twice. Drugs were administered intramuscular (IM) using a cumulative dosing procedure and each drug dose increased the total cumulative dose by one-fourth or one-half log units. For comparison to subsequent vaccine effects, 0.032 mg/kg, IM naltrexone was administered as an acute pretreatment before a single cumulative fentanyl dose-effect test session. This naltrexone has previously shown to produce an approximate 10-fold shift in fentanyl antinociceptive potency in rhesus monkeys (Cornelissen et al., 2018). Human laboratory studies have suggested an approximate 8 to 10-fold potency shift is the minimum clinically effective necessary for naltrexone (Sullivan et al., 2006). Subsequently, the fentanyl-TT conjugate vaccine was administered at weeks 0, 2, 4, 9, 19, and 44 (Figure 1). Fentanyl and oxycodone dose-effect functions were then redetermined over the course of 43 experimental weeks (Figure 1). Drug administration ceased once the cumulative drug dose produced ≥70% rate suppression in individual monkeys.

**Figure 1:**
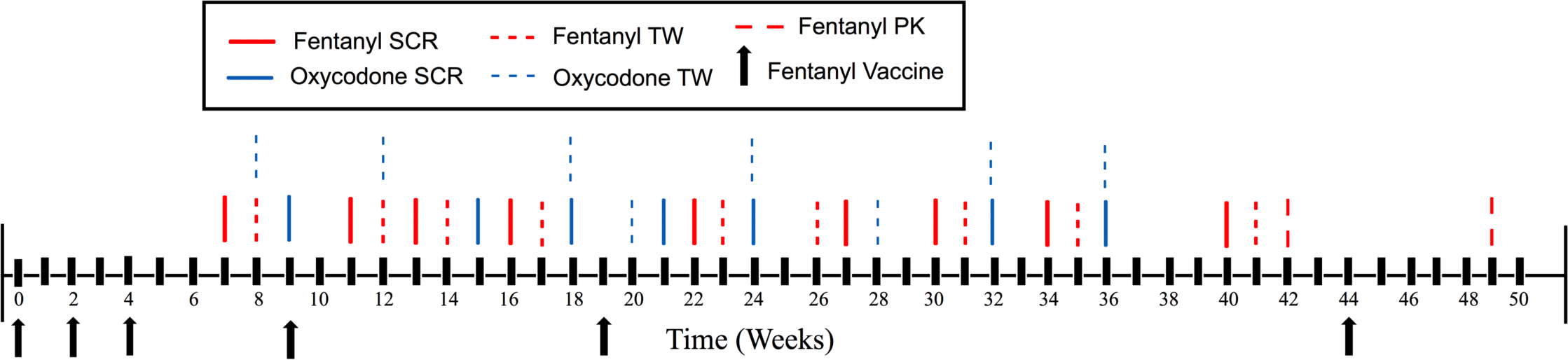
Experimental timeline. SCR stands for schedule-controlled responding; TW stands for tail withdrawal procedure; PK stands for pharmacokinetic experiment.

### 2.3 Thermal nociception procedure

Subjects were also trained to sit comfortably in acrylic restraint chairs as described previously (Banks et al., 2010; Cornelissen et al., 2018). The bottom 10-12 cm of each tail was shaved weekly and immersed in a thermal container of water heated to 38°C or 50°C. Prior to fentanyl and oxycodone dose-effect function determination, a baseline cycle occurred where tail withdrawal latencies must have been 20s at 38°C and ≤2s for 50°C before initiating drug administration. Latencies were recorded with a handheld stopwatch. These criteria were met for each test session. Initially, dose-effect functions were determined for fentanyl (0.001-0.032 mg/kg) and oxycodone (0.01-1.0 mg/kg), and each drug was tested twice. Drugs were administered IM using a cumulative dosing procedure and each drug dose increased the total cumulative dose by one-fourth or one-half log units. Each cycle consisted of two components: a 10-min timeout period and a five-min-test period where tail withdrawal latencies were reassessed at 38 and 50°C. Thermal intensities were presented in random sequence. Fentanyl and oxycodone cumulative dose-effect functions were redetermined over the course of 43 weeks (Figure 1). Drug administration ceased once a subject had maximum latencies of 20s in 50°C. If the subject did not remove its tail before the cutoff time, the experimenter removed the tail.

### 2.4 Pharmacokinetic Study

Blood samples (1–2 mLs) from a saphenous vein were collected in Vacutainer tubes containing 3.0 mg of sodium fluoride and 6.0mg sodium ethylenediaminetetraacetic acid before, and 3, 10, 30, 100, 300 min, and 24 h after 0.018 mg/kg, IM fentanyl administration in monkeys trained to present their leg while seated in custom primate restraint chairs. Fentanyl pharmacokinetic studies were conducted both before fentanyl-TT conjugate vaccine administration number six at week 42 and at week 49 (Figure 1). Samples were immediately centrifuged at 1350g for 10 min. The plasma supernatant was transferred into a labeled storage tube and frozen at −80°C until analyzed. Quantitative analysis of fentanyl was based upon a previously described method (Poklis et al., 2016).

### 2.4 Vaccination Period

The fentanyl vaccine was composed of a fentanyl hapten conjugated to tetanus toxoid (TT) as described previously (Bremer et al., 2016) and solubilized in 50% glycerol and 50% phosphate-buffered saline. Fentanyl copies were 23 per protein, based on conjugation with surrogate bovine serum albumin. On a per monkey basis, 400 μg of conjugate fentanyl-TT hapten was mixed with 600 μg CpG ODN 2006 (Eurofins Genomics, Louisville, KY) and 1 mg Alhydrogel adjuvant 2% (InvivoGen, San Diego, CA) for 30 min and then refrigerated for 24h prior to IM administration at approximately 1.2 ml per monkey. Blood was collected from a saphenous vein into vacutainer tubes every two weeks for subsequent analysis. Titer measurements were obtained by ELISA and fentanyl IC_50_ values were obtained by a competitive SPR assay both using fentanyl-BSA as a coating antigen as previously described (Bremer et al., 2016; Bremer et al., 2017).

### 2.5 Data analysis

For the schedule-controlled responding assay, raw rates of responding (responses/s) from each test cycle were converted to percent of control using the average response rate from the previous training session in each individual monkey. Individual % control rate data were then averaged between monkeys to yield group mean results. For the thermal nociception assay, drug effects were expressed as %Maximum Possible Effect (%MPE). The equation was:

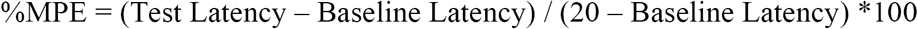

where test latency was tail withdrawal from the 50°C water after drug administration and baseline latency was tail withdrawal at the beginning of the session before drug administration. ED_50_ values were determined for fentanyl and oxycodone in each monkey for each assay as a function of fentanyl vaccine administration. ED_50_ values were calculated by linear regression when at least three data points on the linear portion of the dose-effect function were available or log-linear interpolation (one below and one above 50% effect) as described previously (Bremer et al., 2017; Cornelissen et al., 2018; Cornelissen et al., 2019). Individual ED_50_ values were averaged to yield mean ED_50_s and 95% confidence limits. Potency ratios were calculated by comparing the ED_50_ value during the test condition (vaccine treatment) to the baseline ED_50_ value. Group mean potency ratios were compared using ANOVAs. A Dunnet post-hoc test followed a significant main effect. The criterion for significance was set *a priori* at the 95% confidence level (*p*<0.05).

### 2.6 Drugs and Reagents

Fentanyl HCl, (–)-oxycodone HCl, and (–)-naltrexone HCl were provided by the National Institute on Drug Abuse Drug Supply Program (Bethesda, MD). All drugs were dissolved in sterile water and administered intramuscularly (IM). Drug doses were calculated and expressed using the salt forms listed above.

## 3.0 Results

### 3.1 Vaccine effects on fentanyl and oxycodone-induced rate suppression

Average control rates of responding across all experiments was 1.8 ± 0.1 responses/s. Figure 2 shows the potency of fentanyl (Panel A) and oxycodone (Panel B) to produce rate-suppression before vaccine administration, at week 22 of the experimental timeline (Figure 1) and following acute 0.032 mg/kg naltrexone pretreatment. The corresponding ED_50_ values are reported in Table 1. Acute 0.032 mg/kg naltrexone produced an approximate 13-fold and 8-fold shift in fentanyl and oxycodone rate-suppression potency, respectively. The fentanyl vaccine maximally shifted the fentanyl potency (∼10-fold) at week 22 similar to 0.032 mg/kg naltrexone (Panel A) and in contrast to naltrexone, the vaccine was selective for fentanyl vs. oxycodone (Panel B).

**Table 1:** Potency (ED_50_ value) and 95% confidence limits (CL) of fentanyl and oxycodone to decrease rates of operant responding in male rhesus monkeys (n=4) before and after fentanyl vaccine administration.

| Experimental Week | Fentanyl ED<sub>50</sub> (95% CL) | Oxycodone ED<sub>50</sub> (95% CL) |
| --- | --- | --- |
| Baseline | 0.003 (0.001, 0.006) | 0.05 (0.02, 0.15) |
| + 0.032 mg/kg Naltrexone | 0.049 (0.013, 0.092) | 0.4 (0.1, 1.61) |
| Week 7 | 0.011 (0.001, 0.079) | 0.03 (0.01, 0.11) |
| Week 11 | 0.018 (0.005, 0.068) |  |
| Week 13 | 0.011 (0.002, 0.046) | 0.04 (0.02, 0.08) |
| Week 16 | 0.011 (0.003, 0.047) | 0.05 (0.01, 0.21) |
| Week 22 | 0.028 (0.011, 0.072) | 0.1 (0.05, 0.19) |
| Week 27 | 0.021, (0.006, 0.083) | 0.05 (0.02, 0.18) |
| Week 30 | 0.022 (0.010, 0.045) |  |
| Week 34 | 0.007 (0.002, 0.023) | 0.06 (0.03, 0.12) |
| Week 40 | 0.005 (0.002, 0.017) |  |

**Figure 2:**
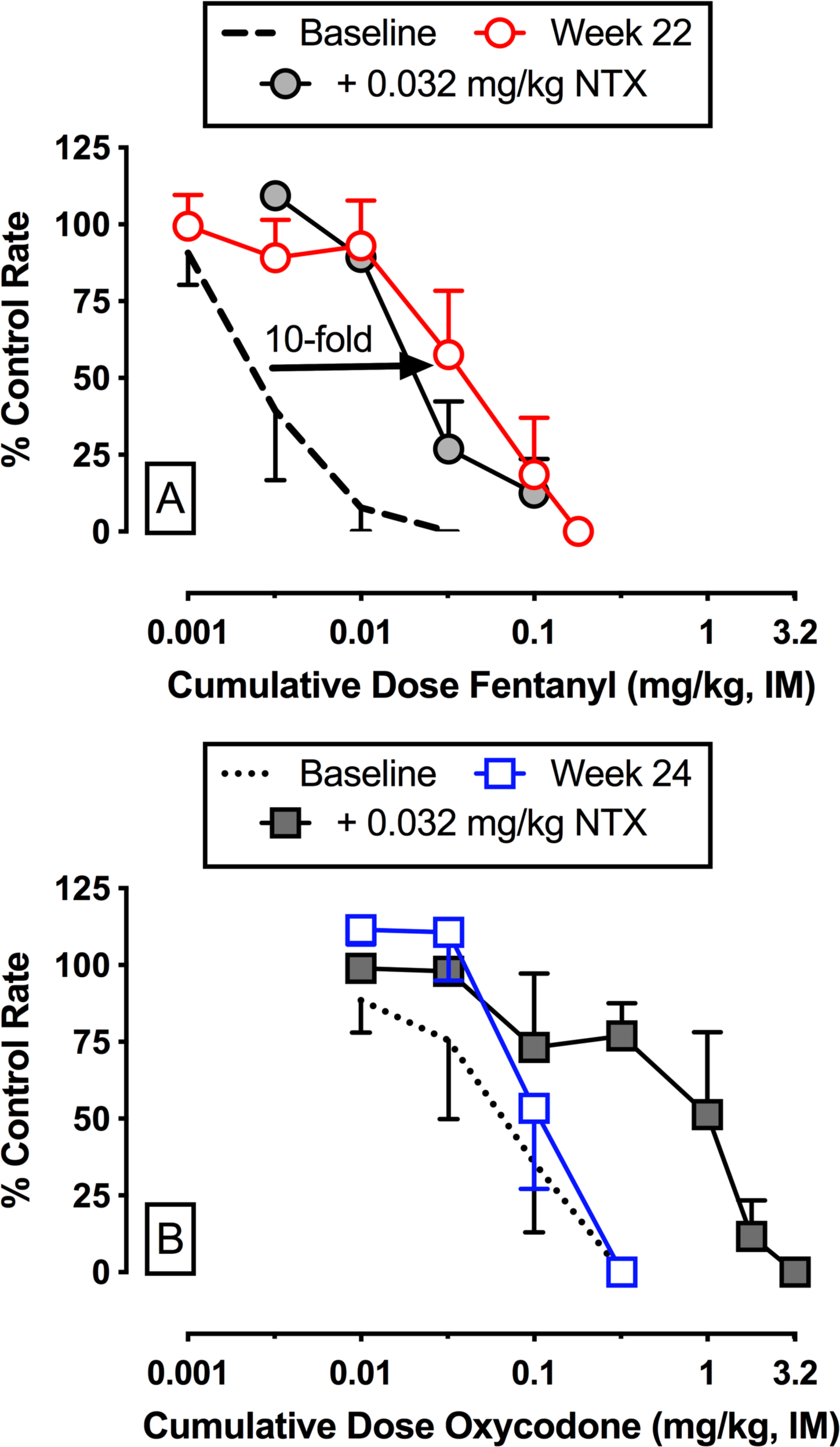
Effects of a fentanyl vaccine and 0.032 mg/kg naltrexone pretreatment on fentanyl (Panel A) and oxycodone (Panel B)-induced rate suppression in male rhesus monkeys (n=4). Abscissae: cumulative intramuscular (IM) drug dose in milligrams per kilogram. Ordinates: percent control rate. All points represent the mean ± SEM from four monkeys. NTX stands for naltrexone.

### 3.2 Vaccine effects on fentanyl and oxycodone-induced antinociception

Average ± S.E.M. baseline tail withdrawal latency before all test sessions was 0.8 ± 0.2 s at 50°C. One monkey that participated in the schedule-controlled responding experiments failed to learn the warm water tail-withdrawal procedure and thus results are from three monkeys. Figure 3 shows the potency of fentanyl (Panel A) and oxycodone (Panel B) to produce antinociception before vaccine administration and at week 12 of the experimental timeline (Figure 1). The corresponding ED_50_ values are reported in Table 2. The fentanyl vaccine maximally shifted the antinociceptive potency of fentanyl 25-fold (Panel A) at week 24 and the vaccine was selective for fentanyl vs. oxycodone (Panel B).

**Table 2:** Antinociceptive potency (ED_50_ value) and 95% confidence limits (CL) of fentanyl and oxycodone in male rhesus monkeys (n=3) before and after fentanyl vaccine administration.

| Experimental Week | Fentanyl ED<sub>50</sub> (95% CL) | Oxycodone ED<sub>50</sub> (95% CL) |
| --- | --- | --- |
| Baseline | 0.004 (0.04, 0.005) | 0.13 (0.06, 0.3) |
| Week 8 | 0.019 (0.003, 0.125) | 0.06 (0.01, 0.25) |
| Week 12 | 0.06 (0.009, 0.4) | 0.13 (0.03, 0.66) |
| Week 14 | 0.06 (0.026, 0.14) | 0.09 (0.04, 0.2) |
| Week 17 | 0.021 (0.008, 0.057) | 0.21 (0.14, 0.32) |
| Week 23 | 0.1 (0.057, 0.175) | 0.43 (0.29, 0.65) |
| Week 26 | 0.06 (0.046, 0.078) | 0.28 (0.13, 0.57) |
| Week 31 | 0.037 (0.018, 0.078) | 0.38 (0.26, 0.57) |
| Week 35 | 0.026 (0.012, 0.055) | 0.31 (0.16, 0.6) |
| Week 41 | 0.024 (0.012, 0.05) |  |

**Figure 3:**
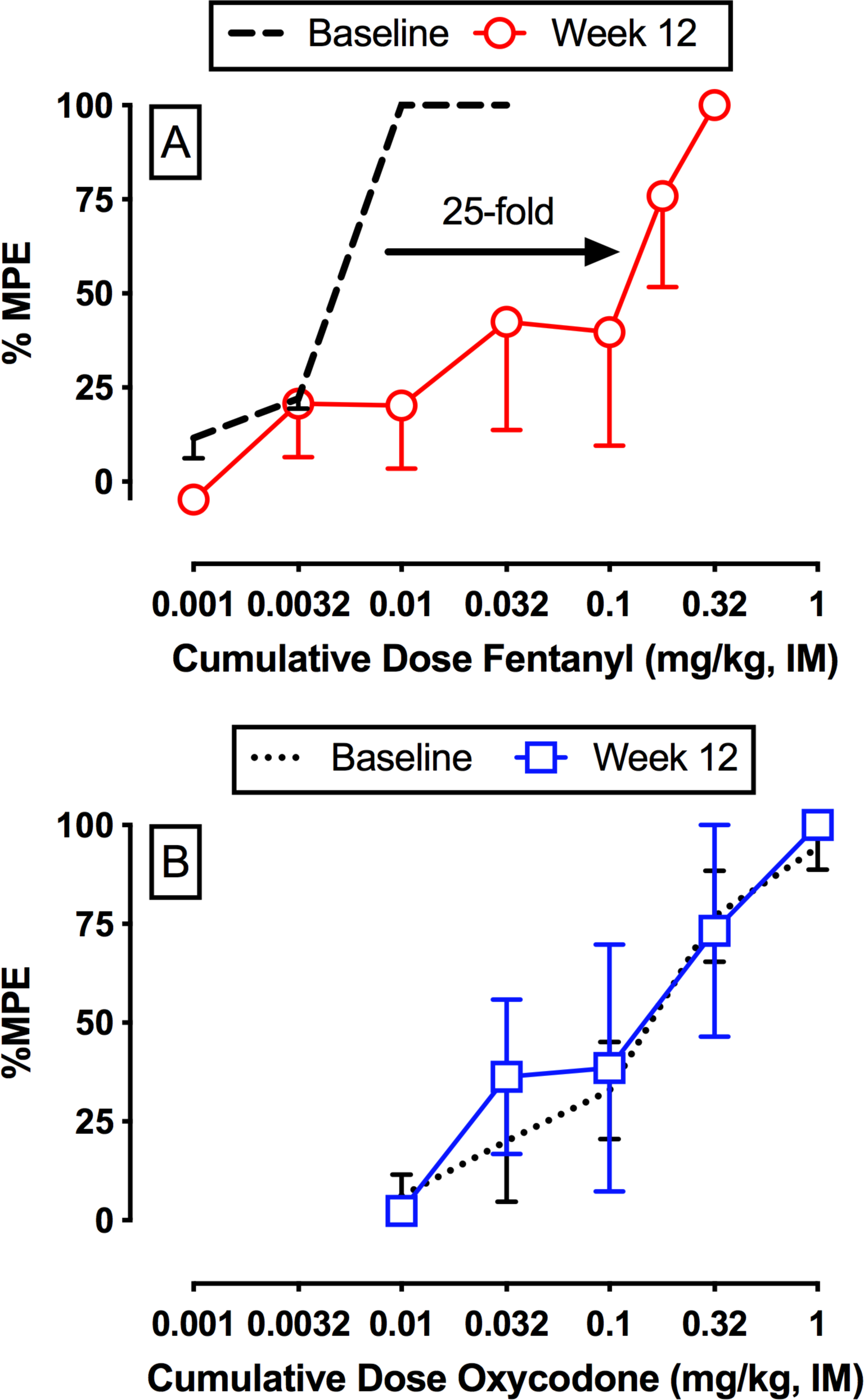
Vaccine effects on fentanyl (Panel A) and oxycodone (Panel B)-induced antinociception in male rhesus monkeys. Abscissae: cumulative intramuscular (IM) drug dose in milligrams per kilogram. Ordinates: percent maximum possible effect (%MPE). All points represent the mean ± SEM from three monkeys.

### 3.3 Time course of fentanyl vaccine effects

Figure 4A shows the changes in fentanyl and oxycodone potency over the entire experimental period in the assay of schedule-controlled responding. The fentanyl vaccine significantly attenuated the fentanyl potency compared to baseline at weeks 11, 22, and 30 (F_9,25.1_=4.15, *p*=0.002) without significantly altering oxycodone potency throughout the entire experimental period. Figure 4B shows the changes in fentanyl and oxycodone potency over the entire experimental period in the warm water tail-withdrawal procedure. The fentanyl vaccine significantly attenuated fentanyl antinociceptive potency compared to baseline at weeks 12 and 23 (F_9,18_=2.48, *p*=0.048). Oxycodone antinociceptive potency was also not significantly altered over the experimental period. Figure 4C shows midpoint titer levels peaked at week 6 and then decayed. Furthermore, midpoint titer levels were positively correlated with fentanyl antinociceptive potency shifts (R^2^=0.28, p=0.0041), but not fentanyl schedule-controlled responding potency shifts (R^2^=0.01). Figure 4D shows antibody-fentanyl affinity (IC_50_ values) maturation over time. Antibody affinity to fentanyl peaked at week 50 (3.2 nM). Moreover, antibody-fentanyl affinity was essentially maximized by week 22 (4.1 nM). Individual midpoint titers and antibody-fentanyl affinities are shown in Supplemental Figure 1.

**Figure 4:**
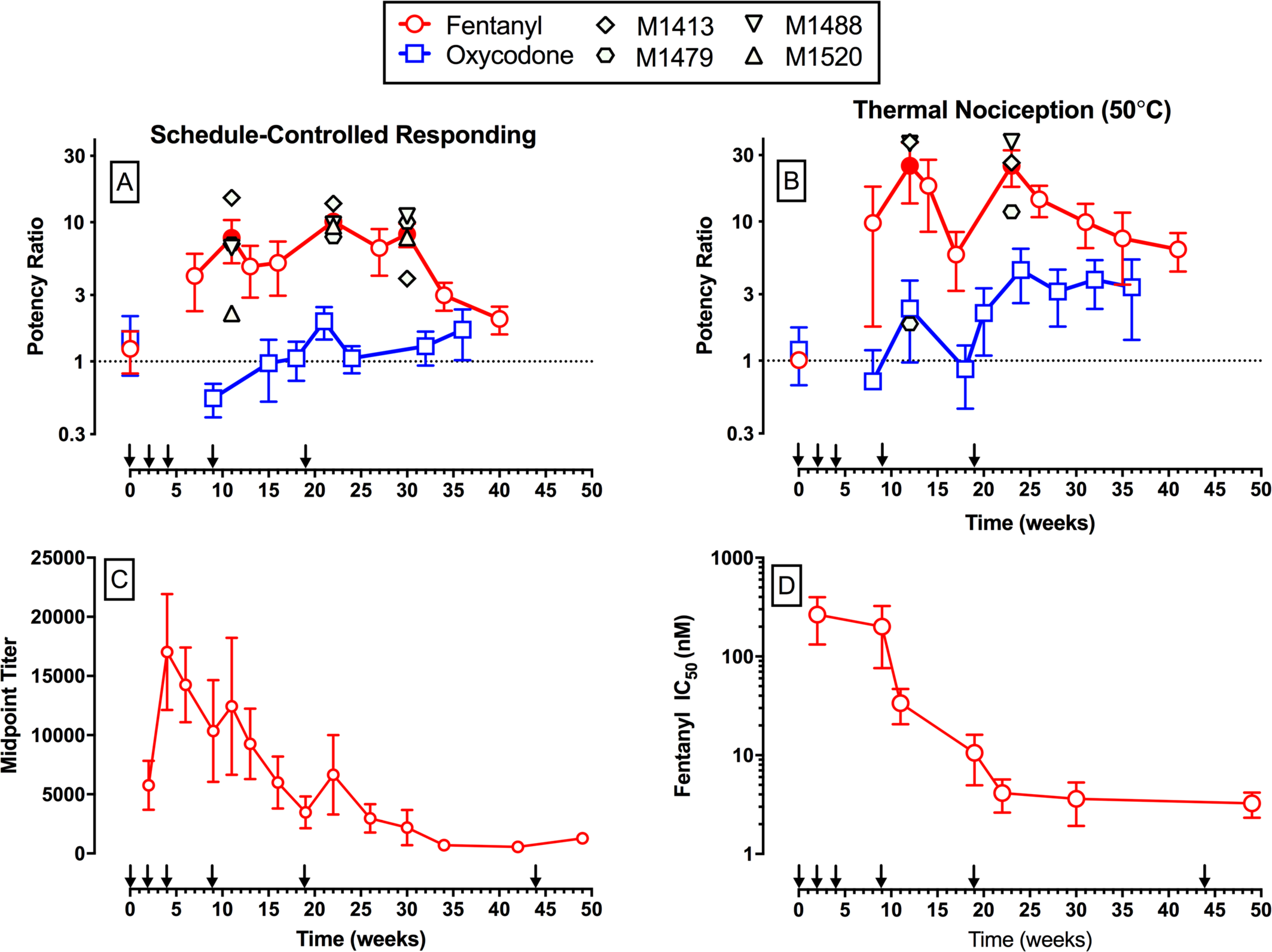
Time course of fentanyl and oxycodone potency shifts in assays of schedule-controlled responding (A) and thermal nociception (B) in male rhesus monkeys. All points represent the mean ± SEM from four monkeys for Panel A and three monkeys for Panel B. Panel C shows midpoint titer levels as a function of experimental week. Panel D shows anti-fentanyl antibody affinity (IC_50_ values, nM) as a function of experimental week. Filled symbols denote statistical significance (*p*<0.05) compared to baseline and individual subject data are shown for each significant data point.

### 3.4 Vaccine effects on fentanyl pharmacokinetics

Figure 5 shows plasma fentanyl levels over time following 0.018 mg/kg fentanyl (IM) administration. Fentanyl levels peaked at 10 min and peak fentanyl levels (41.23 vs. 136 ng/mL) were approximately 3-fold higher 5 weeks following the last fentanyl vaccine boost at week 44. Fentanyl levels were significantly greater following vaccine administration at all time points (time: F_5,15_=12.99, *p*<0.0001; vaccine: F_1,3_=31.67, *p*=0.011; time × vaccine interaction: F_5,15_=5.2, *p*=0.0057).

**Figure 5:**
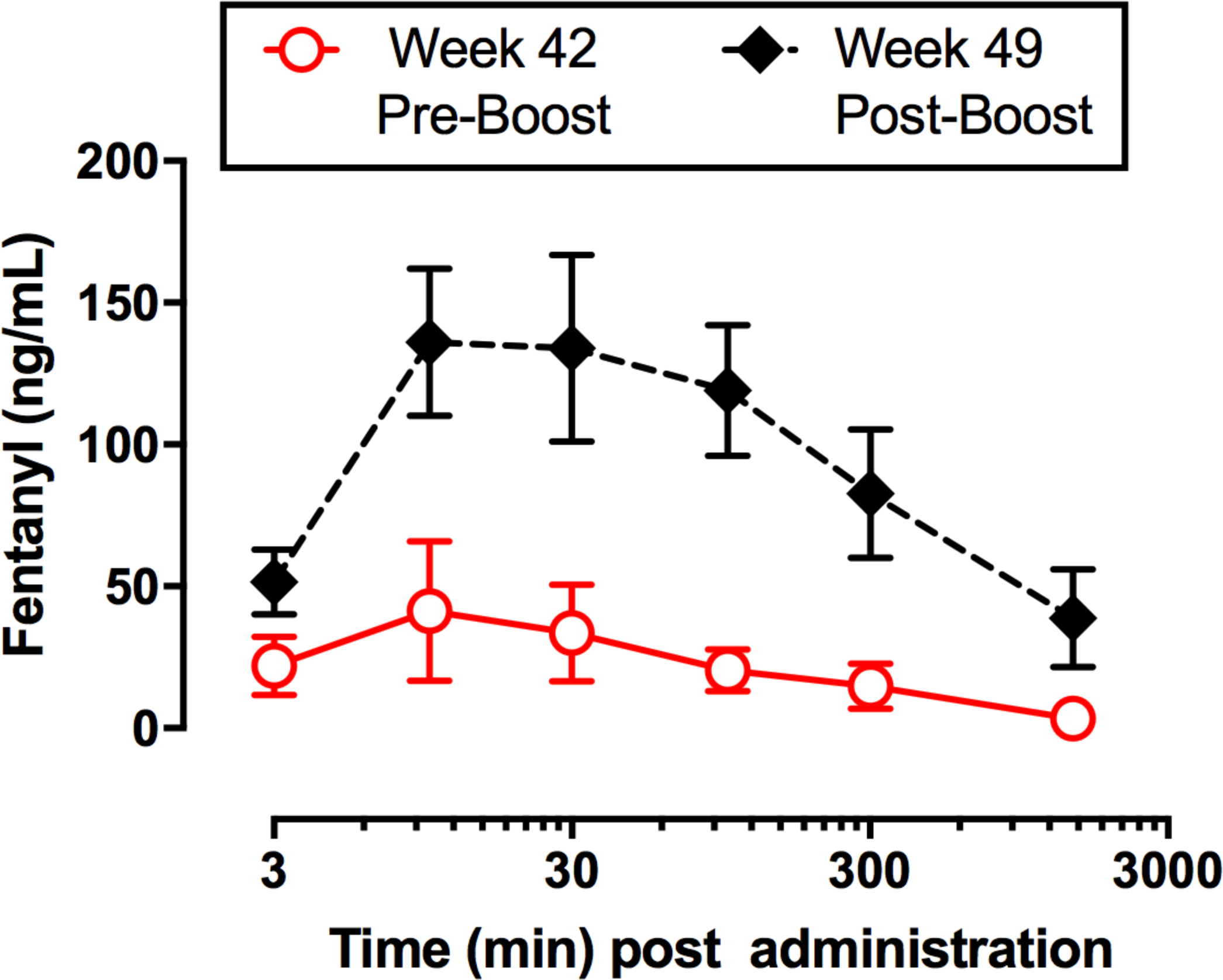
Vaccine effects on fentanyl plasma levels in male rhesus monkeys. All points represent the mean ± SEM from four monkeys. Filled symbols denote statistical significance (*p*<0.05) compared to the week 42 (Pre-Boost).

## 4.0 Discussion

The aim of the present study was to determine the effectiveness and selectivity of a fentanyl-TT conjugate vaccine to alter the behavioral and pharmacokinetics of fentanyl in rhesus monkeys. There were three main findings. First, vaccine administration significantly shifted the potency of fentanyl to produce rate-suppression and antinociception greater than 10-fold. These potency shifts were of similar magnitude to acute 0.032 mg/kg naltrexone administration and the minimum potency shift reported to be necessary for a clinically effective antagonist-based OUD treatment (i.e., depot naltrexone). Second, the fentanyl vaccine was selective for fentanyl compared to the structurally dissimilar MOR agonist oxycodone. Lastly, the vaccine significantly increased plasma fentanyl levels suggesting the antibodies sequestered fentanyl in the blood. Altogether, these results demonstrate that a fentanyl vaccine can produce clinically significant potency shifts in fentanyl behavioral effects and support the continued development and evaluation of this fentanyl vaccine to address the ongoing opioid crisis.

Both fentanyl and oxycodone produced dose-dependent depression of operant behavior and thermal antinociception in rhesus monkeys. The present results are consistent with previous studies in humans (Finch and DeKornfeld, 1967), nonhuman primates (Banks et al., 2010; Maguire and France, 2014; Nussmeier et al., 1991), rodents (Millan, 1989; Schwienteck et al., 2019; Walker et al., 1994). We have previously reported that acute 0.032 mg/kg naltrexone produced an approximate 10-fold potency shift in the fentanyl antinociception dose-effect function at 50°C (Cornelissen et al., 2018). The present results extended these previous acute naltrexone antagonism results of fentanyl to the assay of schedule-controlled responding. Acute 0.032 mg/kg naltrexone produced an approximate 13-fold potency shift in the fentanyl dose-effect function and a 9-fold potency shift in the oxycodone dose-effect function. Human laboratory studies have suggested that an 8-fold potency shift in mu-opioid agonist dose-effect functions is the minimally effective potency shift necessary to produce clinically meaningful effects on opioid use disorder-related endpoints (Comer et al., 2006; Sullivan et al., 2006). Overall, these results with acute naltrexone as a positive control provide an empirical framework for interpretation of subsequent fentanyl vaccine effects.

The fentanyl vaccine was effective and significantly attenuated the potency of fentanyl to depress operant behavior and produce antinociception. The present results in nonhuman primates are consistent with previous fentanyl vaccine effects in both mice and rats (Bremer et al., 2016; Raleigh et al., 2019; Torten et al., 1975). Moreover, the present results extend these previous findings in two ways. First, the maximum fentanyl potency shift ∼25-fold in tail withdrawal) observed in rhesus monkeys was qualitatively similar to the maximal potency shifts (∼33-fold in tail withdrawal) observed in mice with the same fentanyl-TT conjugate vaccine (Bremer et al., 2016) and greater than the maximum potency shift (∼5.4-fold in hot plate) observed in rats with a fentanyl-KLH conjugate vaccine (Raleigh et al., 2019). Second, fentanyl vaccine effectiveness was less in the assay of schedule-controlled responding (∼10-fold) than warm-water tail withdrawal in the same monkeys. Differences in opioid-targeted vaccine effectiveness between schedule-controlled responding in rhesus monkeys and antinociception in mice have also been reported for a heroin-TT conjugate vaccine (Bremer et al., 2017). One potential reason for differential sensitivity of schedule-controlled responding and thermal nociception to immunopharmacotherapies could be related to differences in MOR agonist efficacy requirement. For example, the partial MOR agonist buprenorphine produces near maximal antinociception in male rhesus monkeys at 50°C but fails to significantly depress rates of operant behavior (Cornelissen et al., 2018; Cornelissen et al., 2019). Thus, thermal nociception would be a lower MOR agonist efficacy requiring procedure compared to schedule-controlled responding. Abuse-related endpoints, such as discriminative stimulus effects, are also low MOR agonist efficacy requiring procedures (for review, see (Bergman et al., 2000)). Overall, the present results suggest schedule-controlled responding may provide a more conservative estimate of subsequent clinical immunopharmacotherapy effectiveness than thermal nociception procedures.

In contrast to naltrexone, the fentanyl vaccine selectively attenuated the behavioral effects of fentanyl relative to oxycodone. The present results in rhesus monkeys are consistent with previously reported selectivities of fentanyl vaccines in both mice (Bremer et al., 2016) and rats (Raleigh et al., 2019). One advantage of high antibody specificity by opioid-targeted vaccine would be to maintain the flexibility of a patient being treated with a structurally dissimilar opioid (e.g. oxycodone) for pain management or allow for combination OUD treatments with immunopharmacotherapies and either naltrexone, buprenorphine, or methadone. However, one potential disadvantage of high antibody specificity is a motivated individual could circumvent vaccine effects by misusing a structurally dissimilar opioid. The fentanyl vaccine utilized in the present studies has been shown effectiveness towards a variety of fentanyl analogues, including α-methylfentanyl, 3-methylfentanyl, and carfentanil (Bremer et al., 2016; Hwang et al., 2018b). Yet, this same fentanyl vaccine elicited antibodies displaying very weak affinity towards structurally dissimilar opioids such as heroin (Bremer et al., 2016; Hwang et al., 2018a). Recent preclinical research has explored the development of combination immunopharmacotherapy approaches directed at multiple, structurally dissimilar abused opioids (e.g., fentanyl and heroin) (Hwang et al., 2018b; Hwang et al., 2018c). In conclusion, opioid-targeted vaccines may provide for a unique clinically effective option for OUD treatment.

Four fentanyl vaccine boosts over the course of nine weeks were necessary to produce the initial significant potency shift. Importantly, significant potency shifts were recaptured following a fifth vaccine boost at week 19. The vaccine latency to produce significant shifts in fentanyl potency in the present study suggests that one clinical hurdle to overcome for immunopharmacotherapies is the slow induction phase compared to depot naltrexone or buprenorphine. Immunopharmacotherapy effectiveness depends upon two main factors 1) the production of sufficiently high levels of viable titers by the immunized subject’s immune system and 2) the antibody affinity for the drug. This latter factor is critical because fentanyl’s affinity for the MOR is in the low nanomolar range. Optimizations in hapten copy number or adjuvant and conjugate dosing may shorten the induction time (Bremer et al., 2017; Hwang et al., 2018a). However, the fentanyl vaccine could also be combined with the once a month depot naltrexone (Krupitsky et al., 2011) or buprenorphine (Haight et al., 2019) formations to ensure sufficient antibody titer levels.

In summary, the present results in rhesus monkeys support the continued development of a fentanyl-TT conjugate vaccine to address the opioid crisis. Although the clinical effectiveness of fentanyl or other opioid-targeted vaccines to treat OUD remains to be empirically determined, the clinical deployment of a fentanyl-targeted vaccine could be utilized to address public health harm reduction effects. For example, a fentanyl vaccine could serve as a harm reduction agent to mitigate opioid overdose due to fentanyl or fentanyl analog contaminated heroin (Ciccarone, 2017) or other drugs of abuse (Mars et al., 2018). In addition, a fentanyl vaccine could be utilized to protect service personnel against chemical threats involving fentanyl or fentanyl analogs.

## Acknowledgements

We appreciate the technical assistance of Stacie Havens for the pharmacokinetic experiments.

## Funding Sources and Disclosures

This work was supported by the National Institutes of Health grants (UH3DA041146, P30DA033934, F32AI126628). NIDA had no role in study design, collection, analysis, and interpretation of the data, in the writing or decision to submit the manuscript for publication. The content is solely the responsibility of the authors and does not necessarily represent the official views of NIDA. All authors report no conflicts of interest. KDJ is listed as an inventor on a Scripps Research Institute patent for the fentanyl-TT conjugate vaccine and has been licensed.

