## Supplemental Figure for "Vaccine blunts fentanyl potency in male rhesus monkeys"

**Number of Figures: 1**

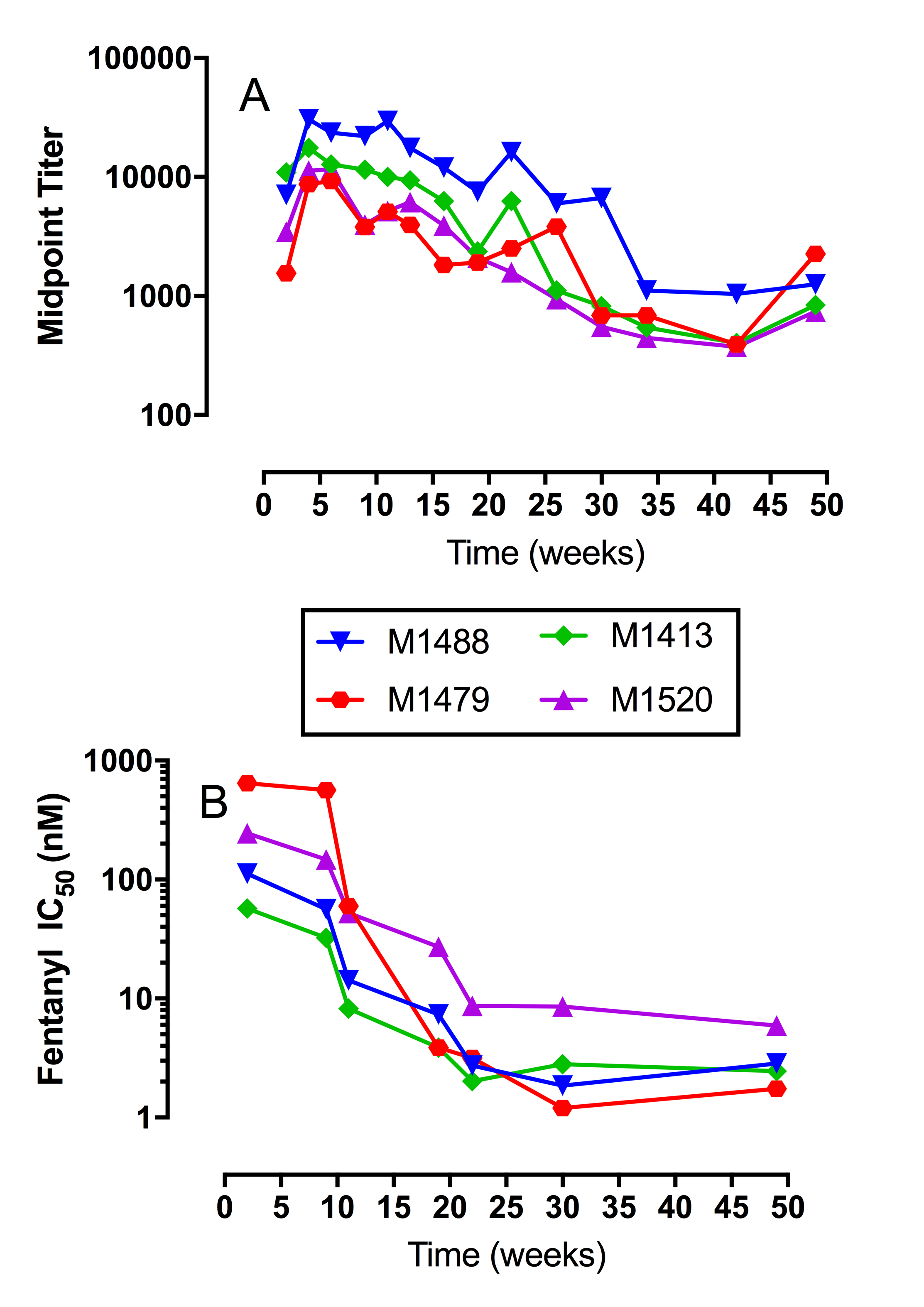

**Figure 1:** Time course of midpoint titers and anti-fentanyl antibody affinities in individual male rhesus monkeys. Panel A shows midpoint titer levels as a function of experimental week. Panel B shows anti-fentanyl antibody affinity (IC_50_ values, nM) as a function of experimental week. All points represent individual subject data.
